# Detecting large-scale networks in the human brain using high-density electroencephalography

**DOI:** 10.1101/077107

**Authors:** Quanying Liu, Seyedehrezvan Farahibozorg, Camillo Porcaro, Nicole Wenderoth, Dante Mantini

**Affiliations:** Neural Control of Movement Laboratory, ETH Zurich, 8057 Zurich, Switzerland; Laboratory of Movement Control and Neuroplasticity, KU Leuven, 3001 Leuven, Belgium; Department of Experimental Psychology, Oxford University, Oxford OX1 3UD, United Kingdom; MRC Cognition and Brain Sciences Unit, Cambridge CB2 7EF, United Kingdom; LET’S-ISTC-CNR, Fatebenefratelli Hospital, 00186 Rome, Italy; Department of Information Engineering, Marche Polytechnic University, 60131 Ancona, Italy.

**Keywords:** electroencephalography, high-density montage, resting state network, functional connectivity, neuronal communication

## Abstract

High-density electroencephalography (hdEEG) is an emerging brain imaging technique that can permit investigating fast dynamics of cortical electrical activity in the healthy and the diseased human brain. Its applications are however currently limited by a number of methodological issues, among which the difficulty in obtaining accurate source localizations. In particular, these issues have so far prevented EEG studies from showing brain networks similar to those previously detected by functional magnetic resonance imaging (fMRI). Here, we report for the first time a robust detection of brain networks from resting state (256-channel) hdEEG recordings. Specifically, we obtained 14 networks previously described in fMRI studies by means of realistic 12-layer head models and eLORETA source localization, together with ICA for functional connectivity analysis. Our analyses revealed three important methodological aspects. First, brain network reconstruction can be improved by performing source localization using the cortex as source space, instead of the whole brain. Second, conducting EEG connectivity analyses in individual space rather than on concatenated datasets may be preferable, as it permits to incorporate realistic information on head modeling and electrode positioning. Third, the use of a wide frequency band leads to an unbiased and generally accurate reconstruction of several network maps, whereas filtering data in a narrow frequency band may enhance the detection of specific networks and penalize that of others. We hope that our methodological work will contribute to rise of hdEEG as a powerful tool for brain research.

## 1. Introduction

Physiological, neuropsychological and neuroimaging studies have clearly revealed that functional specialization and integration are two distinct, yet coexisting principles of human brain organization (Friston, 2002). Specifically, although the function of an area at a given cortical location is highly specialized, the information it processes is dependent on its precise connections with other areas in different parts of the brain (Varela et al., 2001). Large-scale functional interactions between spatially distinct neuronal assemblies can be assessed using functional connectivity methods, which estimate statistical dependence between the dynamic activities of distinct brain areas (Friston, 2011). Functional connectivity is most often measured using functional magnetic resonance imaging (fMRI) data, which have a spatial resolution of a few millimeters and permit to construct accurate maps of large-scale functional networks across the brain (Fox and Raichle, 2007; Ganzetti and Mantini, 2013). However, a significant drawback in the context of functional connectivity is that fMRI provides only an indirect measure of brain activity mediated by a slow hemodynamic response. Alternatively, electroencephalography (EEG) or magnetoencephalography (MEG) can be utilized to estimate large-scale functional interactions within large-scale brain networks. Despite a number of technical limitations, they are potentially more suited to investigating mechanisms of long-range neuronal communication, insofar as they yield high temporal resolution and directly measure electrophysiological activity (Ganzetti and Mantini, 2013; Pfurtscheller and Lopes da Silva, 1999).

In recent years, technological advances have enabled the reliable reconstruction of ongoing activity in the brain (typically called ‘source space’) using MEG (Mantini et al., 2011). These developments have permitted to confirm the electrophysiological basis of fMRI-based connectivity (Brookes et al., 2011; Hipp et al., 2012). For instance, band-limited MEG power across distant brain regions was found to be temporally coherent during rest, and spatially organized similarly to resting state networks (RSNs) previously identified using fMRI (Brookes et al., 2011; de Pasquale et al., 2010). Moreover, MEG studies have begun to disclose important information about brain network dynamics also during task performance (de Pasquale et al., 2012; Hipp et al., 2011), suggesting that long-range neuronal communication is characterized by rapid changes of synchronized oscillatory activity within specific brain circuits (de Pasquale et al., 2010). However, applications of MEG for large-scale studies remain limited, mainly because MEG is not portable and has high maintenance costs.

There may be several reasons why no research group has been able to map brain networks using EEG, as previously done using fMRI (Fox and Raichle, 2007; Ganzetti and Mantini, 2013; Gillebert and Mantini, 2013) and MEG (Brookes et al., 2011; de Pasquale et al., 2010; Hipp et al., 2012). One of the main technical difficulties to obtain RSNs from EEG signals is that the high requirement of accurate and precise source activity reconstructions. Unlike MEG, source analysis of EEG potentials requires indeed precise, realistic biophysical models that incorporate the exact positions of the sensors as well as the properties of head and brain anatomy, such that appropriate source localization techniques can be applied to map surface potentials to cortical sources (Michel et al., 2004). To build a realistic head model, accurate representation of the volume conductor of the head and precise volume conductivity of each tissue are essential (Cho et al., 2015; Fiederer et al., 2015; Haueisen et al., 1997; Ramon et al., 2006). Moreover, spatial sampling density and coverage of EEG electrodes also play a crucial role for neuronal source estimation (Slutzky et al., 2010; Song et al., 2015). High-density EEG (hdEEG) provides both high spatial sampling density and large head coverage, which facilitates the reconstruction of brain activity in the source space. Many research groups working with EEG still make use of low-density systems with 32 or 64 channels, whereas hdEEG systems are not widespread yet. Also, dedicated processing tools that permit to use hdEEG for brain imaging in a manner that is analogous to MEG are currently lacking. Another concern to study RSNs with EEG (also with MEG) is the so-called ‘signal leakage’ across brain voxels (Brookes et al., 2012; Hillebrand et al., 2012; Hipp et al., 2012). In EEG studies, the leakage problem can be caused by volume conduction as well as the ill-posed nature of the inverse solutions. While the former occurs inevitably during the signal recording and at a sensor level, the latter is due to the fact that EEG/MEG source estimation consists of estimating a few thousand voxel activities from maximally a few hundred recordings. Therefore, the source estimation is underspecified in nature and yields a blurred image of the true activity in the brain voxels where activity estimated in one voxel is in fact a weighted sum of the activities in the neighboring voxels.

Here we propose that higher degrees of freedom needed to correctly resolve the dynamics of brain activity can be achieved through increasing the number of sensors by utilizing hdEEG. Furthermore, MEG studies documented that the signal leakage problem is less critical when detecting RSNs with independent component analysis (ICA) than seed-based connectivity analysis (Brookes et al., 2011). ICA performs a blind decomposition of a given number of spatio-temporal patterns that are mixed in the data, assuming that these patterns are mutually and statistically independent in space (sICA) or time (tICA). For fMRI analyses, the use of sICA is warranted because the number of time points is typically much smaller than that of brain voxels, and this possibly leads to unreliable data decomposition by tICA (McKeown et al., 1998). However, tICA has been preferred in EEG/MEG connectivity studies (Brookes et al., 2011; Yuan et al., 2016). In the case of EEG/MEG, the use of tICA is possibly not problematic due to the higher temporal resolution of these techniques as compared to fMRI. No study has ever tested whether and to what extent sICA can successfully retrieve brain networks from EEG/MEG data.

In this study, we describe a complete pipeline for the detection of EEG RSNs, which exploits the advantages of high-density as compared to low-density EEG systems and includes state-of-the-art tools for data preprocessing, realistic head model generation, source localization and ICA-based connectivity analysis. Notably, hdEEG data can be collected simultaneously with fMRI and also in combination with non-invasive brain perturbation by transcranial magnetic stimulation (TMS) or transcranial direct/alternating current stimulation (tDCS/tACS). Furthermore, hdEEG experiments can be easily performed not only in healthy volunteers but also in neurological and psychiatric patients. Our methodological work may therefore open up new exciting research avenues in neuroscience, and contribute to rise of hdEEG as a powerful tool for both basic and translational investigations on human brain networks.

## 2. Materials and Methods

### 2.1 Data collection

Data used in this study comprise resting-state hdEEG signals, electrode positions and individual whole-head anatomy MRI from nineteen healthy right-handed subjects (age 28 ± 5.9 years, 5 males and 14 females). All participants reported normal or corrected-to-normal vision, had no psychiatric or neurological history, were free of psychotropic or vasoactive medication. Before undergoing the examination, they gave their written informed consent to the experimental procedures, which were approved by the local Institutional Ethics Committee of ETH Zurich.

The EEG experiment was performed in accordance with the approved guidelines, in a quiet, air-conditioned laboratory with soft natural light. Continuous 5-minute resting EEG data with eyes open were collected. To reduce eye movements and blinks, subjects were instructed to keep fixation on the center of screen during the experiment. High-density EEG signals were recorded at 1000 Hz by the 256-channel HydroCel Geodesic Sensor Net (GSN) using silver chloride–plated carbon-fiber electrode pellets provided by Electrical Geodesics (EGI, Eugene, Oregon, USA). During recording, the EGI system used the electrode at vertex (labeled as Cz in the 10/20 international system) as physical reference. In addition, to better characterize the scalp distribution of EEG signals, all 256 sensors and three landmarks positions (nasion, left and right preauricolar) were localized prior to the EEG acquisition by using a Geodesic Photogrammetry System (GPS). In detail, GPS derives the position of each EEG electrode from multiple pictures, simultaneously captured, of all the sensors on the subject’s scalp. After defining the 2D electrode positions on at least 2 pictures, 3D coordinates are computed by using a triangulation algorithm (Russell et al., 2005). In addition to EEG data and electrode position information, a T1-weighted whole-head MR image of each subject was acquired in a separate experimental session using a Philips 3T Ingenia scanner with a turbo field echo sequence. The scanning parameters were: TR=8.25ms, TE=3.8ms, 8° flip angle, 240×240×160 field of view, 1 mm isotropic resolution.

### 2.2 Method for EEG network detection

We developed a complete analysis workflow to obtain multiple subject-specific RSNs from hdEEG recordings (Fig.1). Four main analysis steps are involved: 1) *Data preprocessing*, to attenuate noise and artifacts that are mixed in the data; 2) *Volume conduction model creation*, to establish how brain sources (i.e. ionic currents in the cortex) can generate specific distributions of potentials over the hdEEG sensors; 3) *Brain activity reconstruction*, to estimate-based on the EEG recordings and the head model-the distribution of active brain sources that most likely generates the potentials measured over the hdEEG sensors; 4) *Connectivity analysis*, to obtain RSN maps showing brain regions that have similar modulations of power, and are therefore thought to preferentially interact with each other. The software implementing the analysis workflow described above is freely available, and can be found at http://www.bindgroup.eu/index.php/software.

**Fig. 1.**
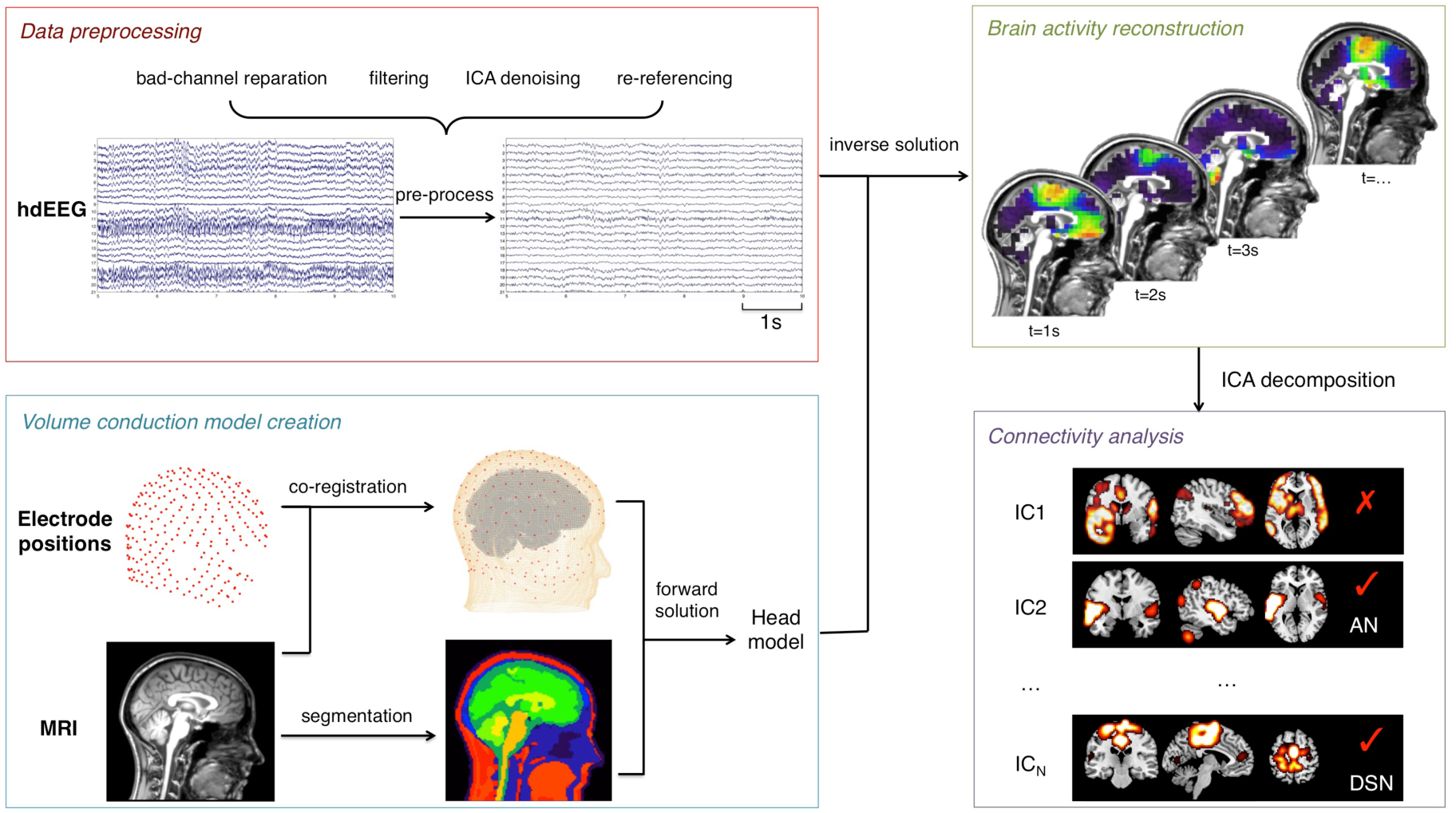
Pipeline for obtaining RSNs from hdEEG recordings. The main analysis steps include: 1) Data preprocessing, involving bad-channel detection, filtering, ICA-denoising and re-referencing; 2) Volume conduction model creation, involving electrodes co-registration, MRI segmentation and forward modeling solution; 3) Brain activity reconstruction, to estimate the distribution of active brain sources that most likely generates the potentials measured over the hdEEG sensors; 4) Connectivity analysis, extracting independent components from the power time series of voxels and selecting the components associated with large-scale brain network activity.

#### 2.2.1 Data preprocessing

First of all, we detected channels with low signal quality and labeled them as ‘bad channels’. To this end, we used an automated procedure that combines information from two different parameters. The first parameter was the minimum Pearson correlation of the signal in a frequency band of interest (here we selected the band 1-80Hz) against all the signals from the other channels. The second parameter was the noise variance, estimated in band in which the contribution of the EEG signal can be considered negligible (here we selected the band 200-250Hz). We define bad channels those for which at least one of the two channel-specific parameters were outliers as compared to the total distribution of values. To ensure robustness of the detection, the threshold to define an outlier was set at equal to *m*+4*s*, where *m* is the average value and *s* is the standard deviation. The detected bad channels were interpolated by using information from the neighboring channels, as implemented in the FieldTrip toolbox (http://www.fieldtriptoolbox.org). Later, we band-pass filtered the data in the frequency range 1-80Hz and we applied independent component analysis (ICA) in order to remove of ocular and muscular artifacts (Mantini et al., 2008). A fast fixed-point ICA (FastICA) algorithm (http://research.ics.aalto.fi/ica/fastica) using a deflation approach and hyperbolic tangent as contrast function was used to extract independent components (ICs). After ICA decomposition, the artifactual ICs were automatically classified by extracting and assessing the following parameters: 1) correlation *c*_*p*_ between the power of the IC with vertical electrooculogram (vEOG), horizontal electrooculogram (hEOG) and electromyogram (EMG) (see Supplementary Fig. 1); 2) the coefficient of determination *r*^*2*^ obtained by fitting the IC power spectrum with a 1/f function; 3) the kurtosis *k* of the IC. An IC was classified as artifactual if at least one of the above parameters was above a given threshold (Supplementary Table 1), which was set in accordance with previous studies (de Pasquale et al., 2010; Mantini et al., 2009). Finally, following artifact rejection we re-referenced the EEG signals using the average reference approach, which showed to be both robust and accurate when using hdEEG data (Liu et al., 2015).

#### 2.2.2 Volume conduction model creation

Precision and accuracy are essential to retrieve the electrical activity origins in the brain. Specifically, obtaining an accurate EEG forward solution requires the generation of realistic volume conductor model from an individual MR image and the definition of correct electrodes locations with respect to it.

Since electrode positions and MR anatomy are not in the same space, we spatially coregistered the EEG electrodes to MR space (Supplementary Fig. 2). This procedure consisted of three distinct steps. In the first step, we estimated the positions of three anatomical landmarks (nasion, left and right preauricolar) in the MR image by projecting the corresponding predefined Montreal Neurological Institute (MNI) coordinates ([0, 85, −30], [−86, −16, −40] and [86, −16, −40]) to individual space. Then, we calculated a rigid-body transformation to match the three landmarks in electrode space to the corresponding landmarks in MR space, and applied it to the electrode positions (Supplementary Fig. 2A). In the second step, we aligned the electrode positions to the surface of the head extracted from individual MR image (Supplementary Fig. 2B) using the Iterative Closest Point (ICP) registration algorithm (Besl and Mckay, 1992). In the third and last step, we ensured that each electrode was perfectly lying over the head surface by projecting it onto the surface point with the smallest Euclidean distance (Supplementary Fig. 2C).

A realistic head model requires the definition of multiple tissue classes of the head, each characterized by a specific conductivity value. We opted for a solution involving 12 tissue classes (skin, eyes, muscle, fat, spongy bone, compact bone, cortical gray matter, cerebellar gray matter, cortical white matter, cerebellar white matter, cerebrospinal fluid and brain stem), which represents the current state-of-the-art for studies modeling the effect of electrical stimulation on the brain (Holdefer et al., 2006; Wagner et al., 2014). This is putatively more accurate than other solutions typically used in EEG analysis, and involving five or less tissue classes (Fuchs et al., 2002; Wolters et al., 2006). Given the intrinsic difficulty in defining all 12-tissue classes directly from the MR image (Supplementary Fig. 3), we warped a high-resolution head template to subject space using the normalization tool in SPM12 (http://www.fil.ion.ucl.ac.uk/spm/software/spm12). This head template was obtained from the ITIS foundation of ETH Zurich (http://www.itis.ethz.ch/virtual-population/regional-human-models/mida-model/mida-v1-0) (Iacono et al., 2015). The conductivity value associated with each tissue class was defined based on relevant literature (Haueisen et al., 1997), and is in line with recent brain stimulation studies (Holdefer et al., 2006) (Supplementary Table 2).

For the numerical approximation of the volume conduction model, we used the whole-head finite element method (FEM) technique. FEM have been proven to be very effective for solving partial differential equations with complicated solution domain and boundary conditions (Wolters et al., 2004). A prerequisite for FEM is the generation of a mesh that represents the geometric and electric properties of the head volume conductor. A hexahedral mesh (i.e. the points of the mesh are connected to create hexahedrons) of the 12 compartments was generated directly from the warped template image. The dipoles corresponding to brain sources were placed on a regular 6-mm grid spanning the cortical grey matter and cerebellar grey matter. In this study, the leadfield matrix L, which contains the measured potentials corresponding to each configuration of dipole position and orientation, was calculated using the Simbio FEM method integrated in FieldTrip. Based on the reciprocity principle, the scalp electric potentials can be expressed in the following equation with leadfield matrix.

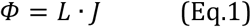

where *L* ∈ ℝ^*N*_*E*_×(3*N*_*v*_)^ is the leadfield matrix; *Φ* ∈ ℝ^*N*_*E*_×1^ is the scalp electric potential; *J* ∈ ℝ^3*N*_*V*_×1^ is the current density at the source; *N*_*E*_ is the number of electrodes, and *N*_*v*_ the number of dipole sources in the cortical grey matter and cerebellar grey matter.

#### 2.2.3 Brain activity reconstruction

We performed reconstruction of brain activity in the source space based on the hdEEG artifact-free recordings and the volume conduction model. To this end, we used the exact low-resolution brain electromagnetic tomography (eLORETA) to perform source reconstruction (Pascual-Marqui et al., 2011). The primary feature of the eLORETA algorithm is that of yielding zero localization error to point sources under ideal (noise-free) conditions. eLORETA estimates the matrix of source activity in the brain J based on the following formula:

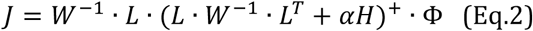

where the superscript + denotes the Moore-Penrose pseudoinverse, *α* >0 is the Tikhonov regularization parameter, *W* ∈ ℝ^*N*_*V*_×*N*_*V*_^ is a symmetric positive definite weight matrix and *H* ∈ ℝ^*N*_*E*_×*N*_*E*_^ is a matrix that depends on the EEG reference. Since the EEG data are in average reference, 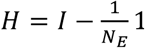, where *I* ∈ ℝ^*N*_*E*_×*N*_*E*_^ is the identity matrix; 1 ∈ ℝ^*N*_*E*_×*N*_*E*_^ where all elements are equal to 1.

The regularization parameter α was estimated by covariance matrix of the noise in measurements, 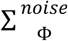, with 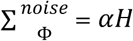. The weight matrix W was iterated until the convergence with 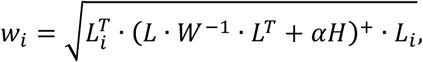 where *w*_*i*_ (i=1, 2,…, *N*_*V*_) is the element of the diagonal weight matrix W.

By estimating the matrix J (see Eq. 2), we obtained the oscillation strength in each dipole with x, y and z orientations at each temporal moment, indicated with j_x_(t), j_y_(t) and j_z_(t) respectively, we obtained the power time series p(t) by means of the following formula:

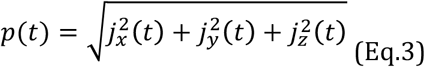

One important issue to measure large-scale connectivity is the effects of signal transmission delays between the distant brain regions (Deco et al., 2011). To avoid the impacts of time delay between long-range sources, we downsampled the power time series to 1 Hz, following an established approach that was proposed in MEG connectivity studies (Brookes et al., 2011). This downsampling can enhance the temporal correlations between brain regions which permits a more accurate detection of slow fluctuation of band-limited power (Supplementary Figs. 4 and 5), and is also well matched with the infra slow fluctuations of the blood oxygen level dependent (BOLD) signal (Palva and Palva, 2012).

#### 2.2.4 Connectivity analysis

Connectivity analysis based on the reconstructed power timecourses across voxels was performed using ICA, in either its spatial or temporal version (Calhoun et al., 2001; Calhoun et al., 2009; Smith et al., 2012). ICA yields a number of independent components (ICs), each of which consists of a spatial map and an associated time-course. The IC spatial map reveals brain regions that have a similar response pattern, and are therefore functional connected (Brookes et al., 2011; Mantini et al., 2007). The number of ICs was estimated by using the minimum description length (MDL) criterion (Li et al., 2007). The FastICA algorithm was run 10 times using a deflation approach and hyperbolic tangent as contrast function to extract reliable ICs, as estimated by the ICASSO software package (Himberg and Hyvarinen, 2003) (http://research.ics.aalto.fi/ica/icasso). EEG-RSNs of interest were selected by using a template-matching procedure. First, the templates were warped to individual MR space, in which the EEG-RSNs were defined. The Pearson correlation was used to estimate the similarity in the spatial distribution of the EEG-ICs and the template RSN maps (Supplementary Fig. 6). The best EEG-IC match for each template map was extracted iteratively, labeled as a specific EEG-RSN, and removed from the pool of EEG-ICs. Accordingly, the same IC could not be associated with two different templates.

### 2.3 Evaluation of brain network reconstruction

First, we applied our hdEEG processing pipeline and reconstructed power envelopes of oscillatory activity in source space from each hdEEG dataset. First of all, we attempted the detection of EEG-RSNs with tICA, following a network detection approach suggested in previous MEG connectivity studies (Brookes et al., 2011). As such, band-limited power envelopes were reconstructed in individual space (whole-brain source space), transformed to common MNI space using SPM and finally concatenated across subjects before running functional connectivity analyses by means of tICA. We then compared this approach with similar approaches in which the source space is constrained to the cortex and/or tICA-based connectivity is run on each single datasets and the resulting network maps are subsequently transformed to MNI space. Power envelopes were separately calculated for the following frequency bands: delta (1-4 Hz), theta (4-8 Hz), alpha (8-13 Hz), beta (13-30 Hz) and gamma (30-80 Hz). We also conducted functional connectivity analyses on power envelopes in a wide frequency range (1-80 Hz), such that the spatial pattern of the reconstructed networks would not be biased by the selection of a specific frequency band. We also examined the possibility of using sICA in alternative to tICA for the detection of EEG brain networks,

In all the analyses described above, we used as templates for RSN detection the maps obtained using from fMRI data used in one of our previous studies (Mantini et al., 2013). These corresponded to: default mode network (DMN), dorsal attention network (DAN), ventral attention network (VAN), right frontoparietal network (rFPN), left frontoparietal network (lFPN), language network (LN), cingulo-opercular network (CON), auditory network (AN), ventral somatomotor network (VSN), dorsal somatomotor network (DSN), visual foveal network (VFN), visual peripheral network (VPN), medial prefrontal network (MPN) and lateral prefrontal network (LPN) (see Supplementary Fig. 6). After the definition of EEG-RSN maps in each subject, we derived a group-level RSN map by using performing a voxel-wise non-parametric permutation test by FSL (http://fsl.fmrib.ox.ac.uk/fsl/fslwiki). We used 5000 permutations for this across-subject analysis, and we set the significance threshold to p<0.01 corrected for multiple comparisons by using the threshold-free cluster enhancement (TFCE) method (Smith and Nichols, 2009).

## 3. Results

First, we attempted to obtain EEG brain networks by applying tICA to alpha-band power envelopes using a whole-brain grid as source space and concatenated datasets, following an approach previous employed in MEG studies. We then compared the results with those obtained when using the cortex instead of the brain as source space, and individual instead of concatenated datasets for network detection (Fig. 2). We specifically evaluated the reconstruction performances focused on the DMN, which presents a complex spatial pattern and is undoubtedly very difficult to reconstruct. Notably, the use of a whole-brain grid as source space resulted in blurred spatial patterns (Fig. 2A,C), whereas all the main DMN areas could be detected only by using a cortex-constrained grid and non-concatenated datasets (Fig. 2D). We therefore retained this solution for further analyses.

**Fig. 2.**
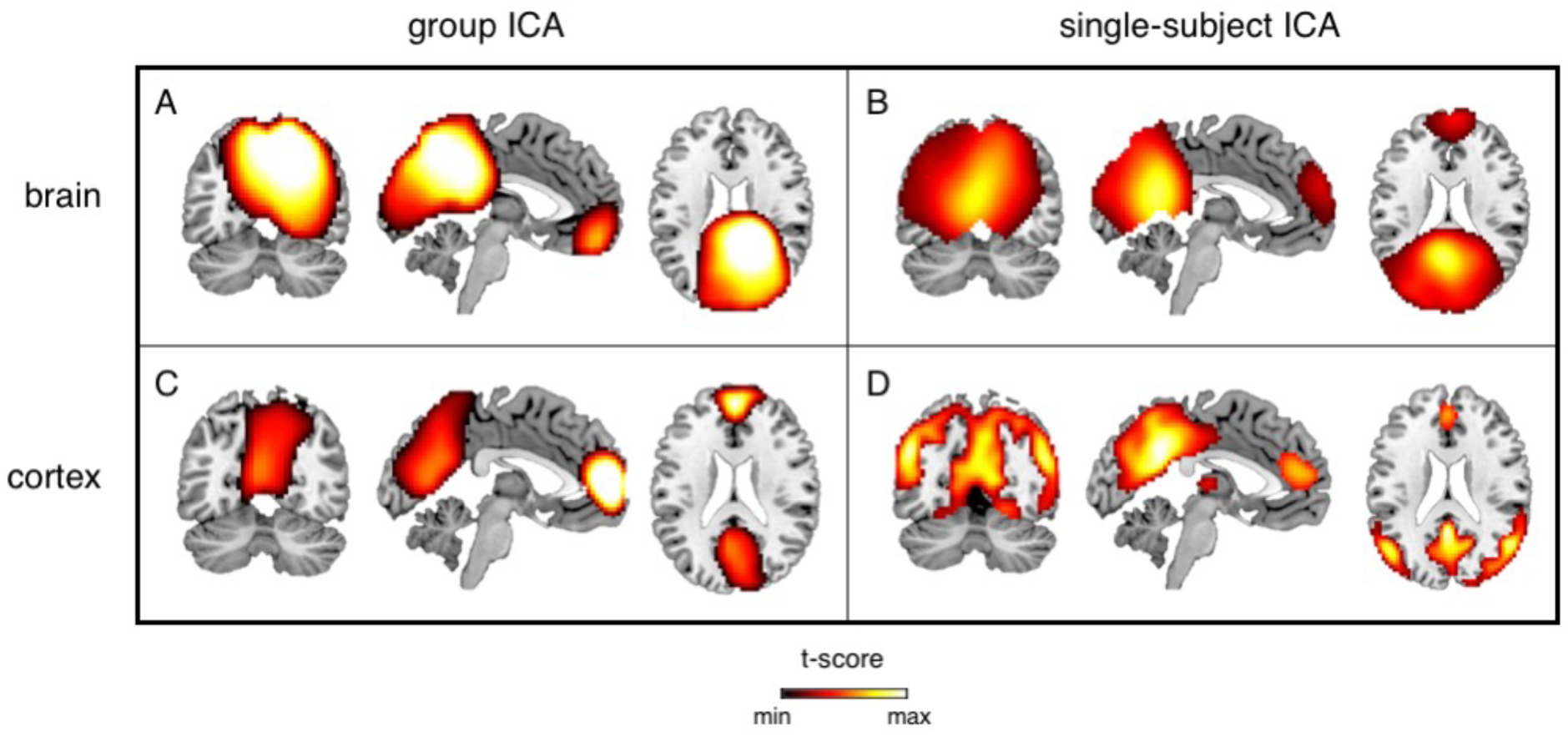
DMN maps reconstructed by tICA using alpha-band power envelopes. (A) Neuronal activity in alpha band (8-13Hz) was estimated from hdEEG data using the whole brain as source space, and brain networks were defined by tICA using concatenated datasets in MNI space. (B) Brain networks were also obtained by using individual datasets (and in individual space). The resulting maps were then transformed to MNI space and subjected to group-level statistical testing. (C-D) The same analysis as in (A) and (B), respectively, was conducted with the cortex as source space instead of the whole brain. Group-level RSN maps (N=19) were thresholded at p<0.01 TFCE-corrected.

There is a general consensus in the literature that neuronal oscillations supporting functional interactions between distant brain regions are mainly in the alpha and beta bands. To test this directly on our data, we examined the performance in RSN reconstruction for different frequency bands. Notably, the DMN could be fully reconstructed using alpha-band power envelopes, and only partially using power envelopes for delta, theta, beta and gamma bands (Fig. 3). When we extended this analysis to other networks (see Methods), we noticed that some of these could be better reconstructed with power envelopes of other frequency bands than the alpha one. There was therefore no frequency band that was optimal for all networks (Supplementary Fig. 7).

**Fig. 3.**
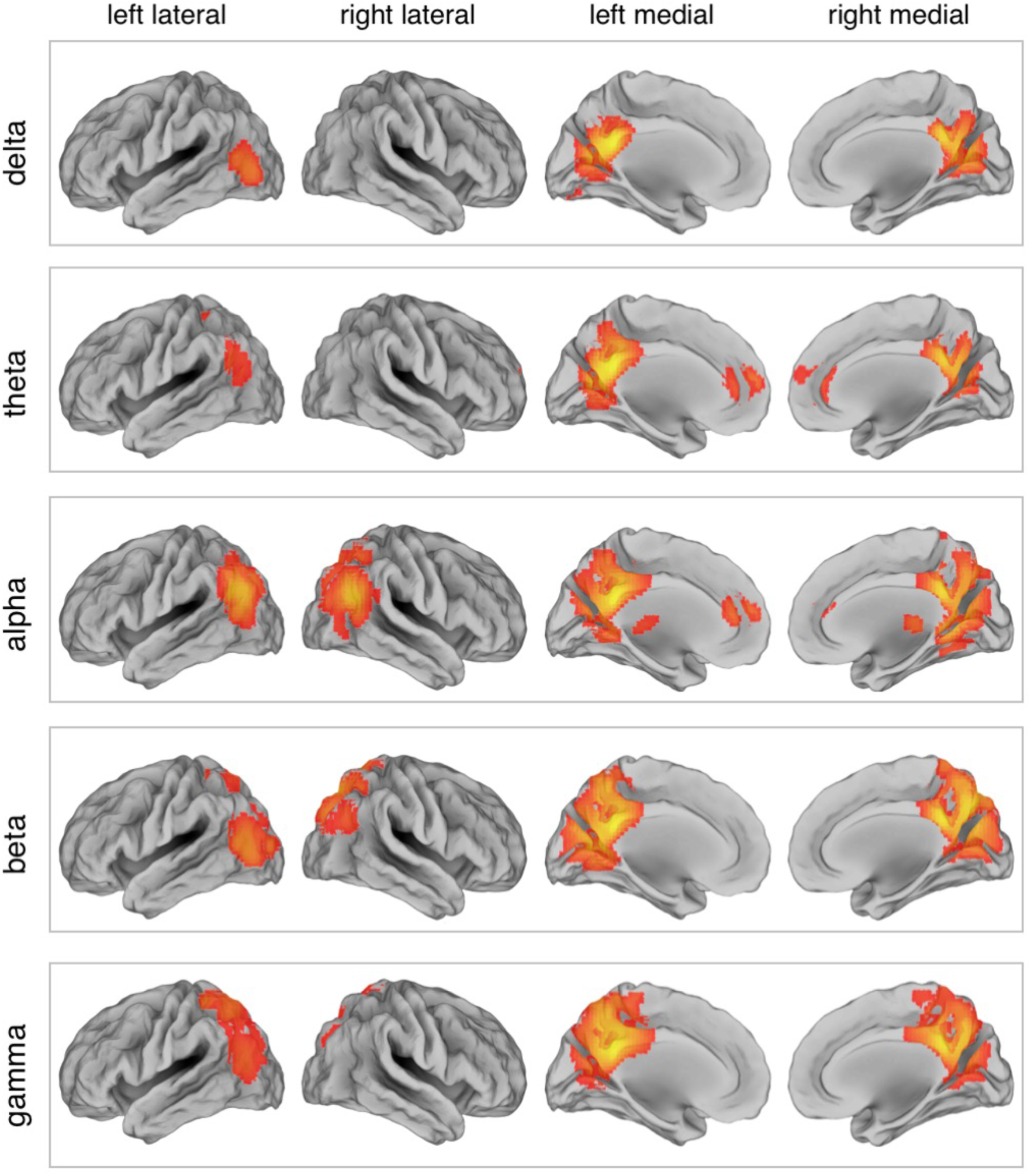
DMN maps obtained by tICA for different frequency bands. The DMN was reconstructed using power envelopes in the delta (1-4 Hz), theta (4-8 Hz), alpha (8-13 Hz), beta (13-30 Hz) and gamma (30-80 Hz) bands, respectively. Surface maps are presented in left/right lateral and medial view (N=19, threshold p<0.01 TFCE-corrected).

We attempted RSN detection by tICA also using EEG signals in a wide frequency band (1-80 Hz). Quite surprisingly, we found in this case that the spatial pattern of each EEG-RSN clearly matched that of the corresponding fMRI-RSN that was used as spatial template (Fig. 4). RSN detection was likewise performed using sICA, always with wide-band EEG signals. Also in this case EEG RSN detection was successful for all networks under investigation (Fig. 5). The EEG-RSN maps obtained with sICA had overall a similar spatial pattern but different features as compared to those obtained by tICA (Supplementary Fig. 8). Specifically, they were less widespread, and covered 40% of the total cortical space, as compared to the 77.8% of those obtained by tICA. The spatial overlap between maps was also smaller with sICA than with tICA, and equal to 4.6% and 39.1%, respectively.

**Fig. 4.**
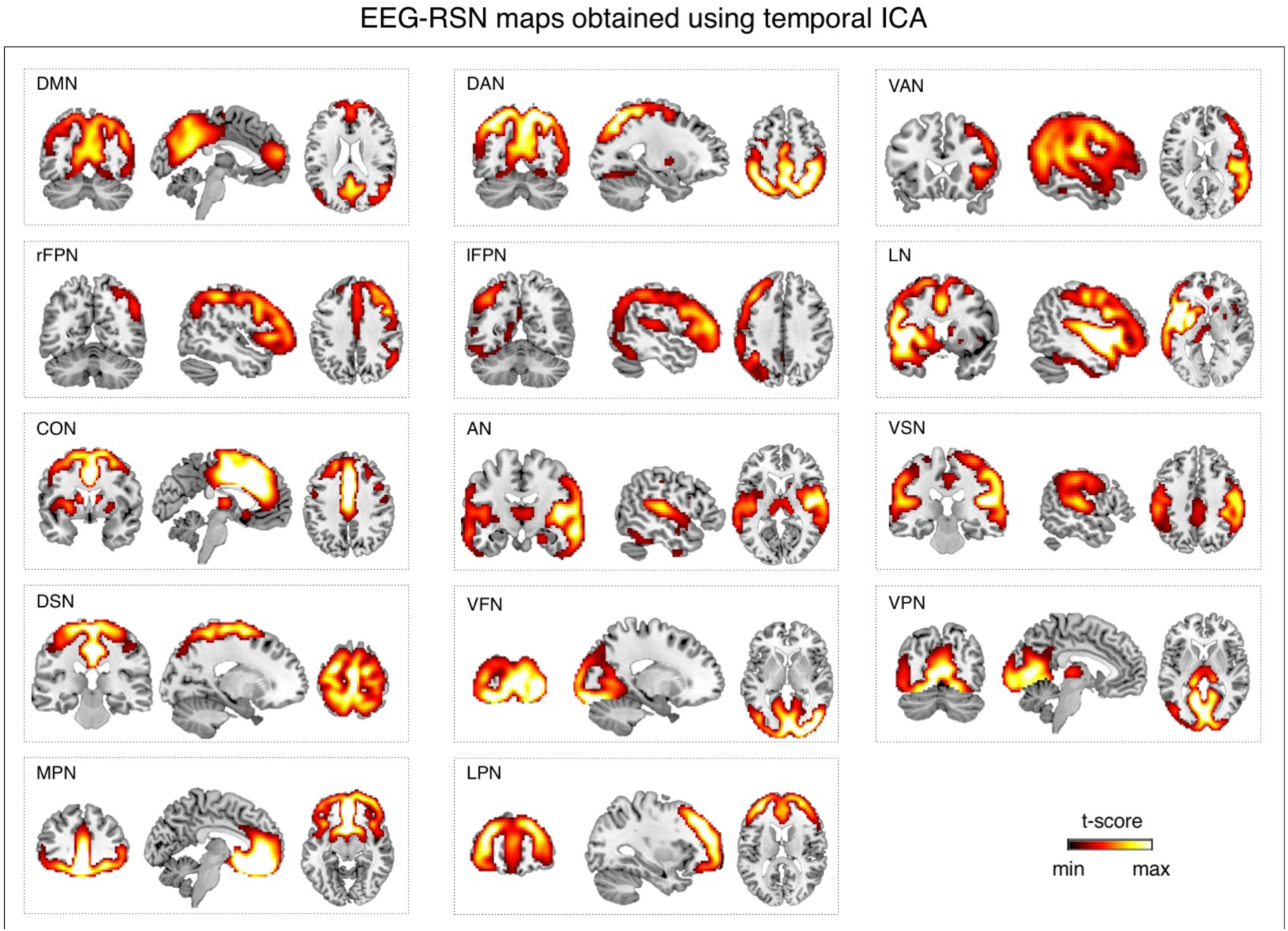
Large-scale brain networks reconstructed using tICA from wide-band EEG signals. EEG networks were selected and labeled on the basis of the spatial overlap with fMRI networks: default mode network (DMN), dorsal attention network (DAN), ventral attention network (VAN), right frontoparietal network (rFPN), left frontoparietal network (lFPN), language network (LN), cingulo-opercular network (CON), auditory network (AN), ventral somatomotor network (VSN), dorsal somatomotor network (DSN), visual foveal network (VFN), visual peripheral network (VPN), medial prefrontal network (MPN) and lateral prefrontal network (LPN). Group-level RSN maps (N=19) were thresholded at p < 0.01 TFCE-corrected.

## 4. Discussion

The main goal of this study was the detection of large-scale brain networks from hdEEG data, with a spatial accuracy comparable to the one that can be obtained using fMRI. This is a particularly complex task, as it requires the precise estimation of neuronal activity in the cortex from recordings made over the scalp. To achieve that goal, we devised a processing pipeline that is tailored to hdEEG data and includes state-of-the-art analysis techniques such as appropriate data pre-processing, realistic head model construction, accurate source localization and ICA-based connectivity analysis. To the best of our knowledge, only one EEG study attempted to reconstruct brain networks using tICA and failed to show maps that correspond to fMRI networks (Yuan et al., 2016). Furthermore, sICA has been extensively used for network detection from fMRI data, but never from EEG/MEG data. Notably, our study revealed that both tICA and sICA can be effectively used for the detection of EEG-RSNs.

### 4.1 Source-space analysis of high-density EEG signals

In this study, we investigated the RSNs spatial patterns using hdEEG. Specifically, we integrated information from hdEEG data, realistic electrode positions and structural MR images. A number of previous studies examined functional connectivity with EEG signals (Smit et al., 2008); however, connectivity analyses were kept at the sensor level due to the low-density electrode coverage. Interpretation of the results of these studies is not straightforward, since EEG recordings contain a mix of neuronal activity from different brain regions. More recently, interest of the scientific community is shifting from low-density EEG toward high-density EEG, from sensor space analyses toward source space analyses, thanks to the technological development and the advanced computing capacity of computers. Our work contributed to the development of analysis tools specifically tailored to hdEEG, providing a novel way to investigate brain activity in a non-invasive manner, and with relatively accurate spatial and temporal resolution.

Previous studies suggested that the use of a realistic head model is essential for retrieving EEG sources (Ramon et al., 2006) and for conducting connectivity analyses in the source space (Cho et al., 2015). In particular, the head model is used to find the scalp potentials that would result from hypothetical dipoles, or more generally from a current distribution inside the head. Accordingly, we paid particular attention to the construction of a realistic head model. First, we used a structural MR image for each subject rather than a template, which was used to achieve a detailed segmentation of the head tissues. A large number of previous studies modeled the head with three compartments, i.e. skull, skin and brain (Fuchs et al., 2002), or five compartments, i.e. skull, skin, white matter, gray matter and cerebrospinal fluid (Van Uitert et al., 2003). Arguing against this oversimplification, we used a finite element model (FEM) based on 12 tissues, following the approach suggested by recent studies that modeled the effect of transcranial electrical stimulation of the brain (Holdefer et al., 2006; Wagner et al., 2014). Moreover, we used electrode positions measured just before the EEG experiment, properly aligned to the segmented MR volume, for the creation of the FEM. Previous work clearly showed the importance of accurate information on electrode positions is crucial for accurate EEG re-referencing (Liu et al., 2015) and source localization (Van Hoey et al., 2000; Wang and Gotman, 2001). However, this is still neglected in a considerable part of current EEG studies.

In our pipeline, we used a realistic head model in combination with eLORETA for the reconstruction of ongoing brain activity in the source space. It should be considered that there is no general consensus about the best EEG source localization method and one should select the method that delivers the best compromise depending on the questions of the study and the data at hand (Michel et al., 2004). Unlike the classical L2 minimum norm estimate (Dale et al., 2000; Lin et al., 2006), eLORETA does suffer from depth bias (Pascual-Marqui et al., 2011). This is an important feature, considering that several crucial nodes in RSNs span deeper brain regions. Furthermore, eLORETA is specifically designed to be minimally affected by the volume conduction error in the EEG (Pascual-Marqui et al., 2011). Whereas resting state MEG studies frequently utilized beamformers for source localization (Baker et al., 2014; Brookes et al., 2011), the suitability of the method for EEG is not established yet. In addition, many MEG studies performed the reconstruction of neuronal activity by using a whole-brain grid for as solution space for source localization (Brookes et al., 2011; Hipp et al., 2012; Marzetti et al., 2013). Our results suggested that having a source space that is confined to the cortex, instead of spanning the whole brain, can improve the reconstruction of brain activity, and therefore of brain networks from high-density EEG data (Fig. 2D). Notably, a cortical constraint to the solution space can be justified from a biophysical point of view, as pyramidal neurons in the cortical gray matter are the principal EEG generators (Schaul, 1998).

### 4.2 Network detection by ICA of power envelopes

We detected RSNs using ICA rather than alternative methods based on seed-based connectivity (Brookes et al., 2012; de Pasquale et al., 2010; de Pasquale et al., 2012), as ICA is a data-driven technique that can produce multiple RSNs by only imposing the constraint of either spatial or temporal independence between RSNs (sICA and tICA, respectively). sICA has been largely employed for the detection of RSNs with fMRI data, in which the number of time points is always much smaller than the number of voxels. In the case of EEG/MEG connectivity studies, tICA has been preferred to sICA since it is possibly better suited to capture the non-linear and non-stationary nature of neurophysiological signals (Brookes et al., 2011; Yuan et al., 2016). Previous MEG connectivity studies performed tICA on band-limited power envelopes, primarily focusing on alpha and beta bands (Brookes et al., 2011). Our connectivity analyses of alpha but also beta power envelopes permitted the robust detection of many, but not all RSNs under investigation (Fig. 3 and Supplementary Fig. 7). Interestingly, we could obtain satisfactory RSN detection when using wideband (1-80 Hz) signals (Figs. 4 and 5). Based on this finding, we argue that the narrow band-pass filtering may not be strictly needed for connectivity analyses. Each brain network is characterized by a specific combination of different neuronal oscillations (Mantini et al., 2007), and the selection of a frequency band may therefore favor the detection of specific networks against others.

**Fig. 5.**
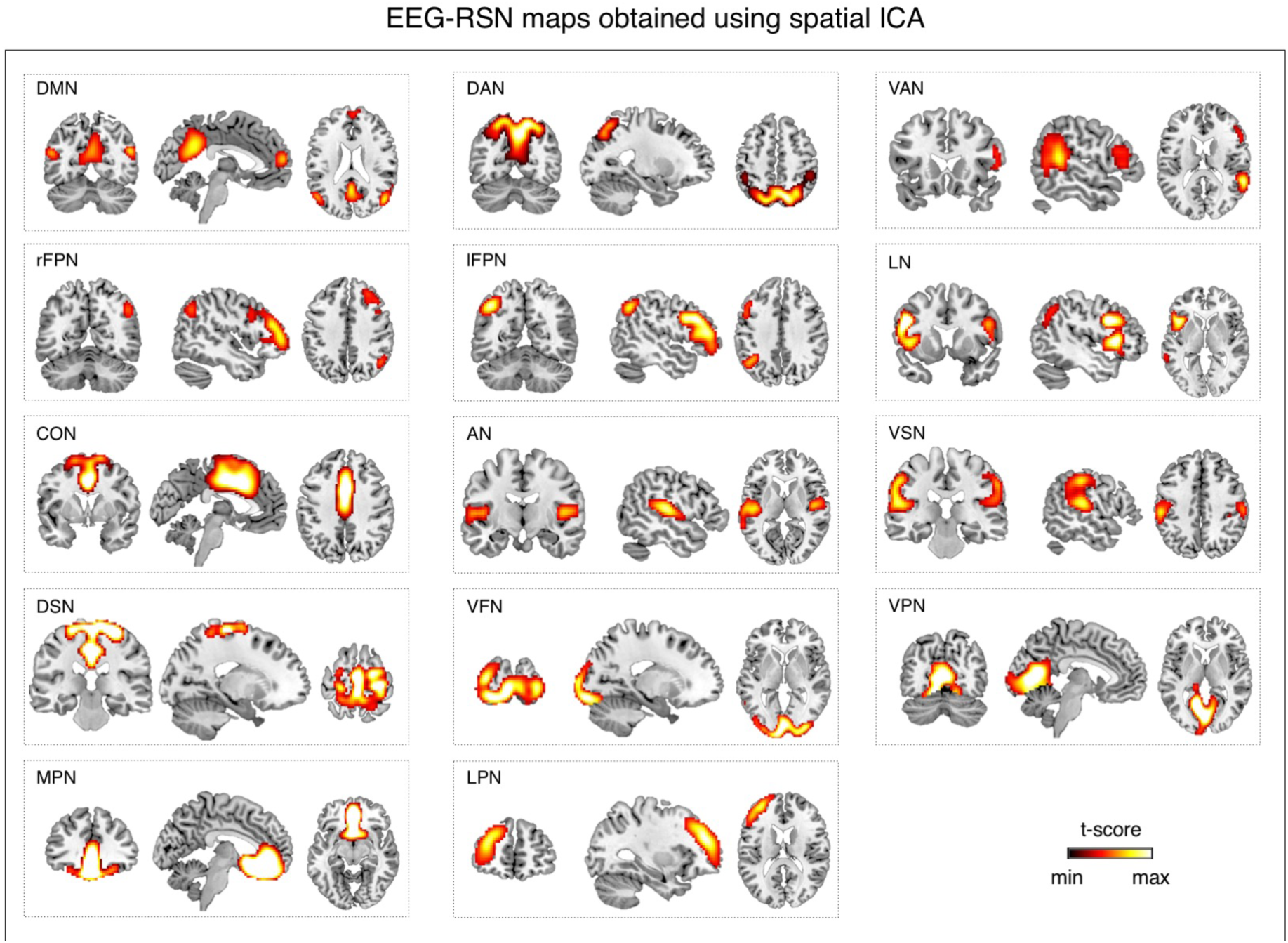
Large-scale brain networks reconstructed using sICA from wide-band EEG signals. EEG networks were selected and labeled on the basis of the spatial overlap with fMRI networks: default mode network (DMN), dorsal attention network (DAN), ventral attention network (VAN), right frontoparietal network (rFPN), left frontoparietal network (lFPN), language network (LN), cingulo-opercular network (CON), auditory network (AN), ventral somatomotor network (VSN), dorsal somatomotor network (DSN), visual foveal network (VFN), visual peripheral network (VPN), medial prefrontal network (MPN) and lateral prefrontal network (LPN). Group-level RSN maps (N=19) were thresholded at p < 0.01 TFCE-corrected.

Overall, our study reveals that both tICA and sICA can be successfully applied for the detection of RSNs from hdEEG data. However, specific differences between RSN maps obtained by tICA and sICA exist. In particular, RSNs with more widespread, sometimes overlapping regions can be observed with tICA, whereas RSNs reconstructed by sICA show more selective spatial patterns and cover more limited portions of the cortical space (Figs. 4 and 5). Overall, our study has the particular merit of showing RSNs that were previously reported only using fMRI but not MEG/EEG, such as VAN, AN and MPN (Mantini et al., 2013). A possible explanation for an increased sensitivity in RSN detection may be the fact that we extracted EEG-RSN maps at the single-subject level and in individual space (Fig. 2D), rather than transforming the source-space power time-courses to common space and performing a single ICA on concatenated time-courses from all subjects (Fig. 2B). The primary reason for our choice is methodological, as this approach permits to better incorporate information on head modeling and electrode positioning (Marino, 2016) in source activity reconstructions. However, it should also be considered that the extraction of RSNs at the single subject level may be important for clinical applications, and in particular for the study of stroke, multiple sclerosis, Alzheimer’s disease and all other conditions in which brain plasticity (Johnston, 2004) may occur.

### 4.3 Study limitations and caveats

The pipeline for the analysis of hdEEG data includes several analysis steps. The successful detection of EEG-RSNs indirectly confirms that each of these steps yielded satisfactory results. From the methodological point of view, an important advancement was the creation of a realistic head model with 12 distinct compartments, which permits to better account for potential spatial distortions in the flow of currents from sources to sensors. It should be noted, however, that our head model did not consider tissue anisotropy. Considering anisotropy may lead to even more accurate results, in particular for subcortical regions (Cho et al., 2015; Wolters et al., 2006). Also, we used conductivity values derived from the literature. A potential improvement may come from the in-vivo estimation of head tissue conductivity, for which techniques are being developed (Akalin Acar et al., 2016; Lew et al., 2009) and may be available in the future. Another potential limitation of the present study pertains to the use of ICA for network detection. Specifically, the number of ICs extracted from the EEG power timecourses was performed using the MDL approach, in line with previous fMRI-RSN studies (Li et al., 2007). Of note, we did not examine how the use of different IC numbers impacts on the quality of the detected RSNs. Future studies are warranted to evaluate if EEG-RSN detection can be further improved by using alternative approaches to estimate the number of ICs.

## 5. Conclusion

In this study, we successfully detected large-scale brain networks using hdEEG data, based on a robust methodology for noise and artifact reduction, head modeling and source localization. The development of such methodology may have broader impact on the field of brain imaging and neuroscience. We posit that hdEEG can constitute a powerful tool for investigating temporal and spectral signatures of long-range functional connectivity in health and disease. Notably, the characterization of functional connectivity dynamics using fMRI is problematic, given the relatively low temporal resolution of the technique. In contrast, EEG – as well as MEG-permits examining network reconfiguration at very fast time scale (Baker et al., 2014; Van de Ville et al., 2010). Moreover, the combination of hdEEG with simultaneous fMRI can unravel the direct relationship between functional connectivity measured through electrophysiological and hemodynamic techniques (Mantini et al., 2007). Finally, analyses of functional connectivity based on hdEEG data may be particularly relevant in a clinical context. In particular, the use of functional connectivity measures from hdEEG has the potential to provide novel and more sensitive biomarkers to improve diagnostics.

## Acknowledgements

The authors would like to thank Joshua Balsters and Marc Bächinger for providing structural MR images based on which head models were built and also Zhiliang Long for scientific discussion about the similarities between EEG and fMRI networks. The work was supported by the Chinese Scholarship Council (scholarship 201306180008 to QL), the Swiss National Science Foundation (grants 320030_146531 and P1EZP3_165207), the Seventh Framework Programme European Commission (grant PCIG12-334039), the KU Leuven Special Research Fund (grant C16/15/070), and the Research Foundation Flanders (FWO) (grants G0F76.16N and G0936.16N).

## Supplementary materials

**Supplementary Fig. 1.**
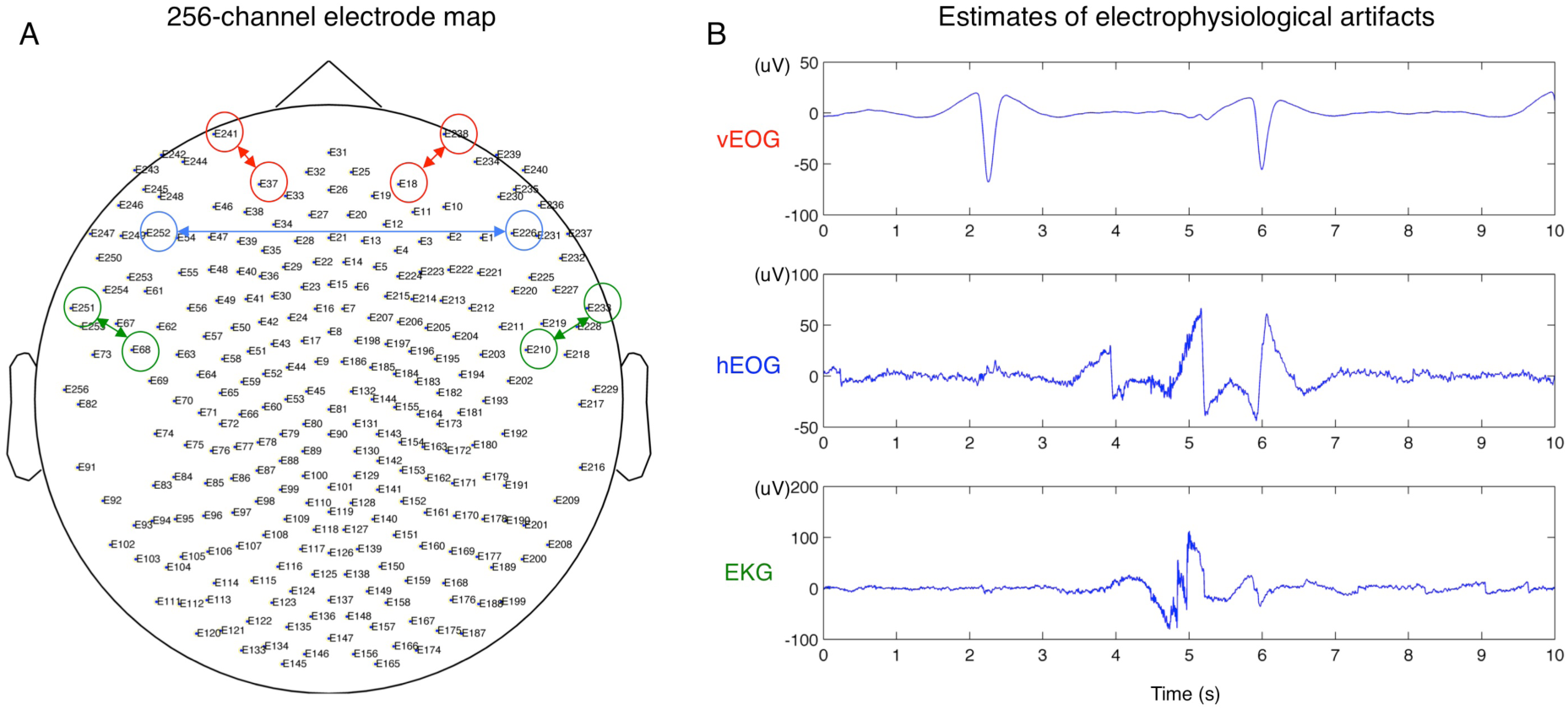
Example of bio-electrophysiological noise signals. By linearly combining EEG signals collected from a 256-channel system (A) we obtained the vertical electrooculogram (vEOG), the horizontal electrooculogram (hEOG) and the electromyogram (EMG) (B).

**Supplementary Fig. 2.**
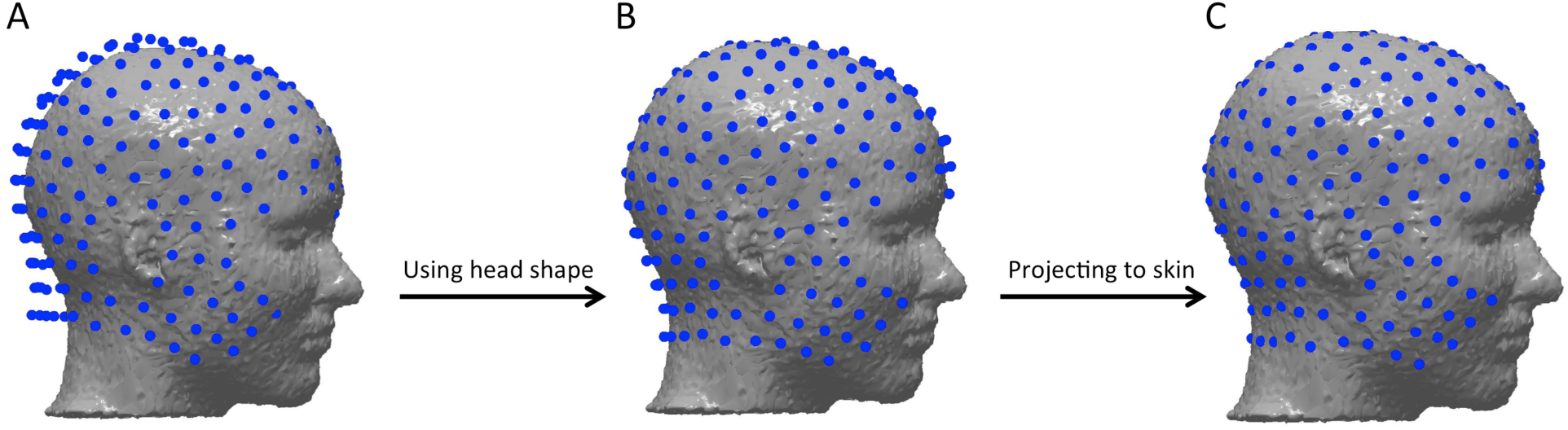
Electrodes co-registration with individual MR image. (A) matching the three landmarks in electrode space with the three landmarks in individual MRI space; (B) using a rigid transformation to match the head shape extracted from the structural MR image with the shape of EEG sensors; (C) projecting the electrodes onto the surface of the head.

**Supplementary Fig. 3.**
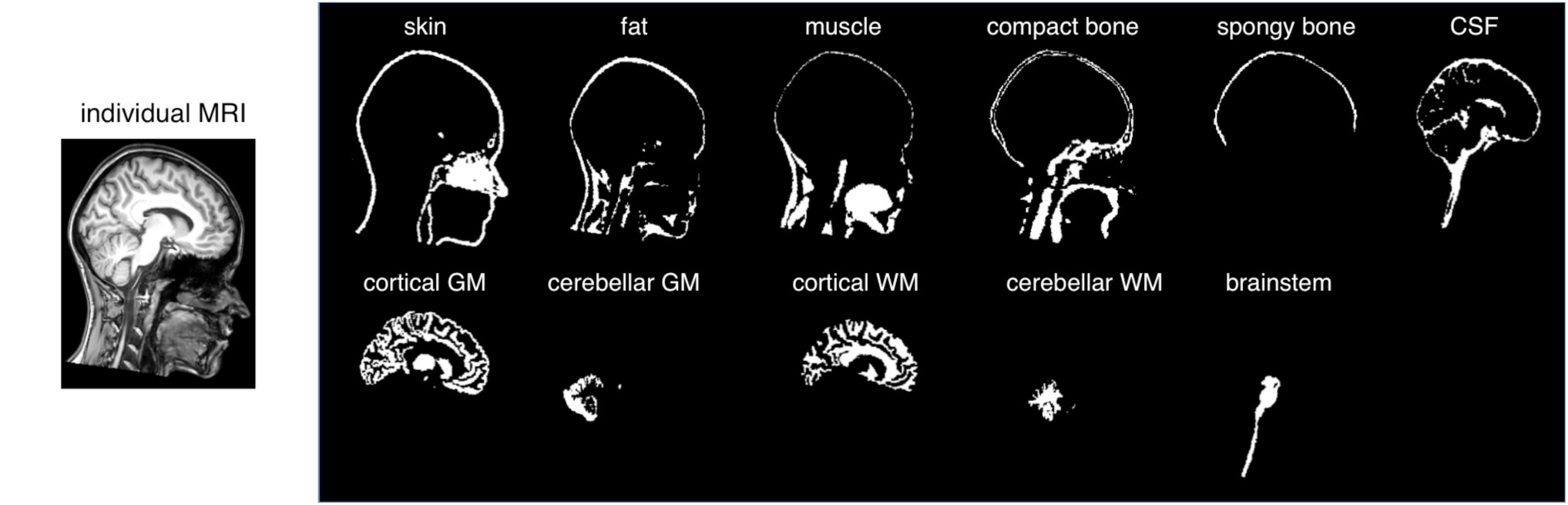
Example of head tissue segmentation using a template warping approach. The MR image of the subject’s head is segmented in 12 compartments: skin, fat, muscle, compact bone, spongy bone, cerebrospinal fluid (CSF), cortical gray matter (GM), cerebellar gray matter, cortical white matter (WM), cerebellar white matter, brain stem, and eyes. An individual MR image is shown in the sagittal section, along with the segmented compartments. Note that eyes are not shown, because they are not visible in the selected MR slice.

**Supplementary Fig. 4.**
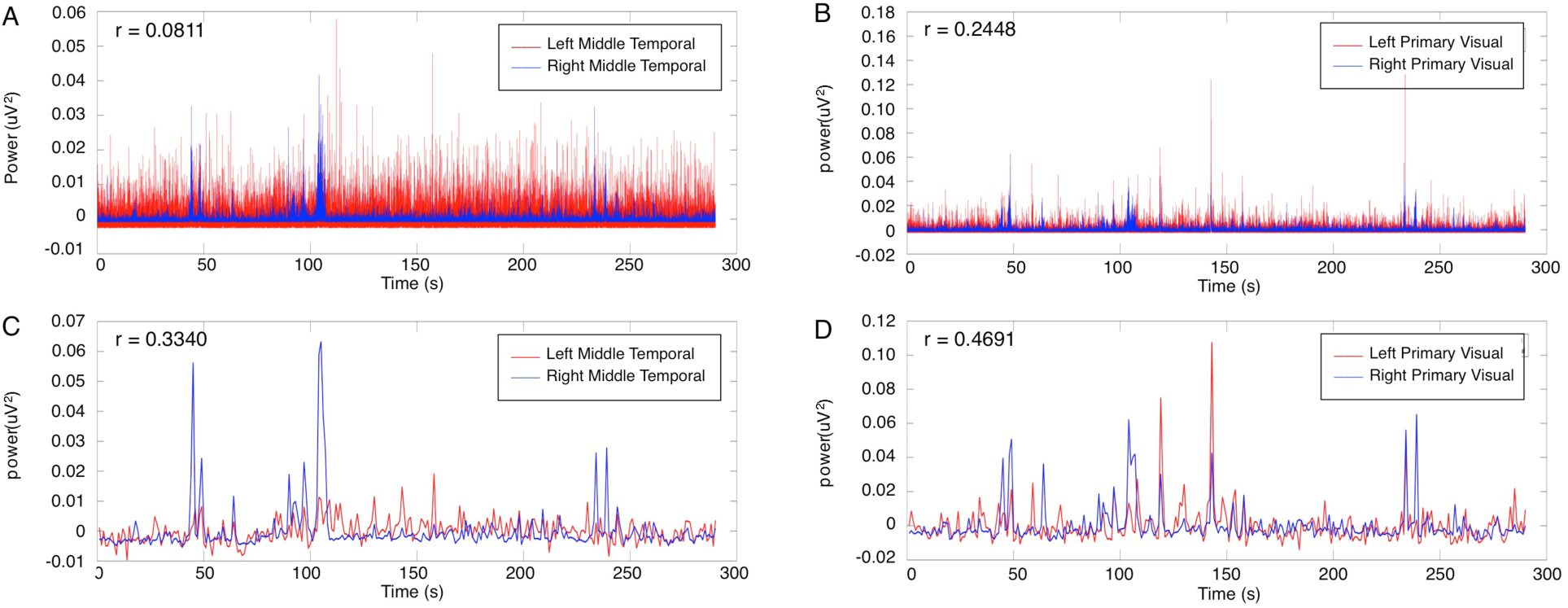
Effect of power envelope downsampling on connectivity detection. We extracted power time series in wideband (1-80Hz) for brain voxels (A) in the left and right middle temporal area, respectively (MNI coordinates: [−43, −72, −8] and [42, −70, −11]), and (B) in the left and right primary visual area respectively (MNI coordinates: [−3, −101, −1] and [11, −88, −4]). We then downsampled the same power time series to 1 Hz (C-D), and examined the temporal correlation between homologous areas before and after downsampling. Notably, this procedure allowed the detection of connectivity that was not observable from the original data.

**Supplementary Fig. 5.**
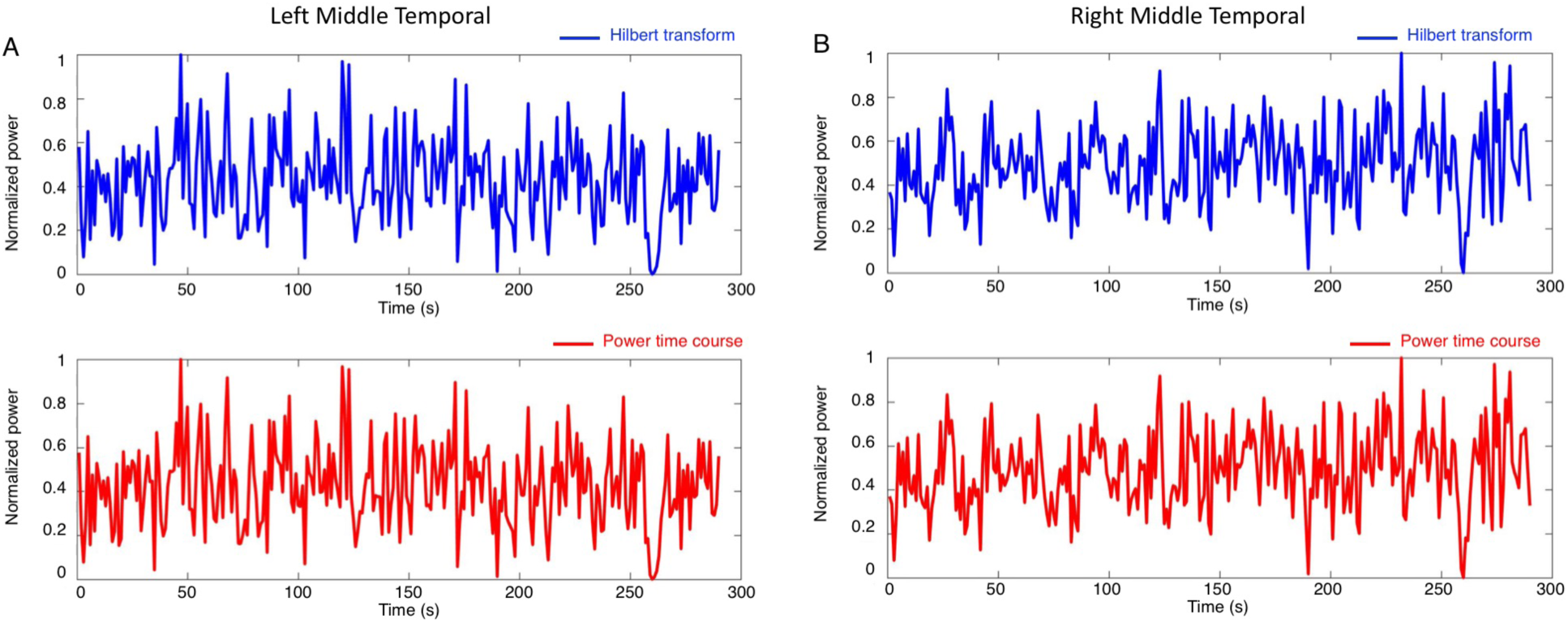
Comparison of different methods to estimate power envelopes. We extracted neuronal activity in alpha band (8-13Hz) in the left (A) and right (B) middle temporal area, respectively (MNI coordinates: [−43, −72, −8] and [42, −70, −11]). We calculated power envelopes by using the Hilbert transform used in Brookes et al. (2011) (blue) and the moving average approach used in de Pasquale et al. (2010) (red). The correlation between the two estimates for the left and right middle temporal area was equal to 0.995 and 0.997, respectively.

**Supplementary Fig. 6.**
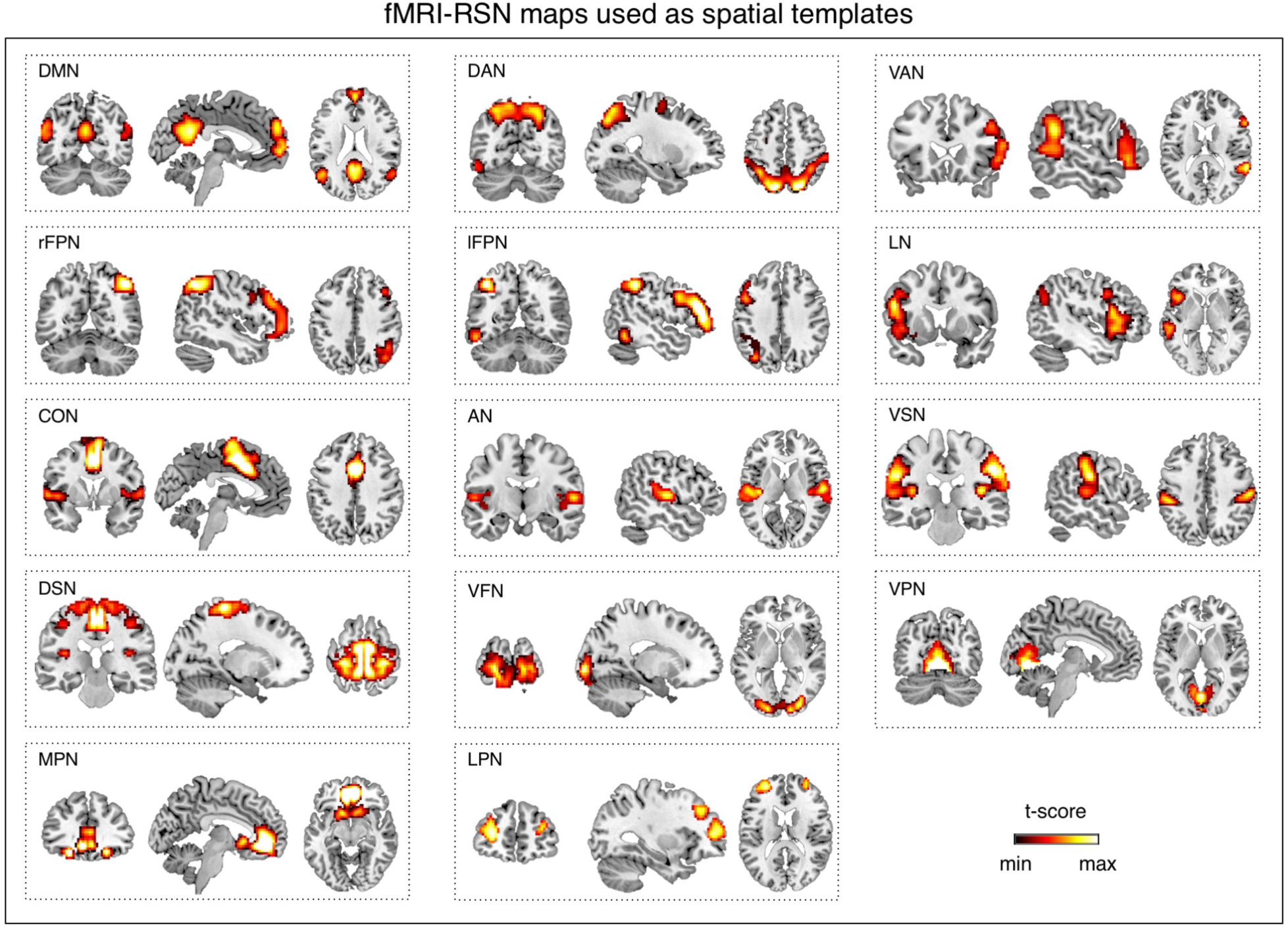
Fourteen fMRI-RSNs maps used as templates in this study. The maps were obtained from twenty-four healthy subjects at rest (Mantini et al., 2013).

**Supplementary Fig. 7.**
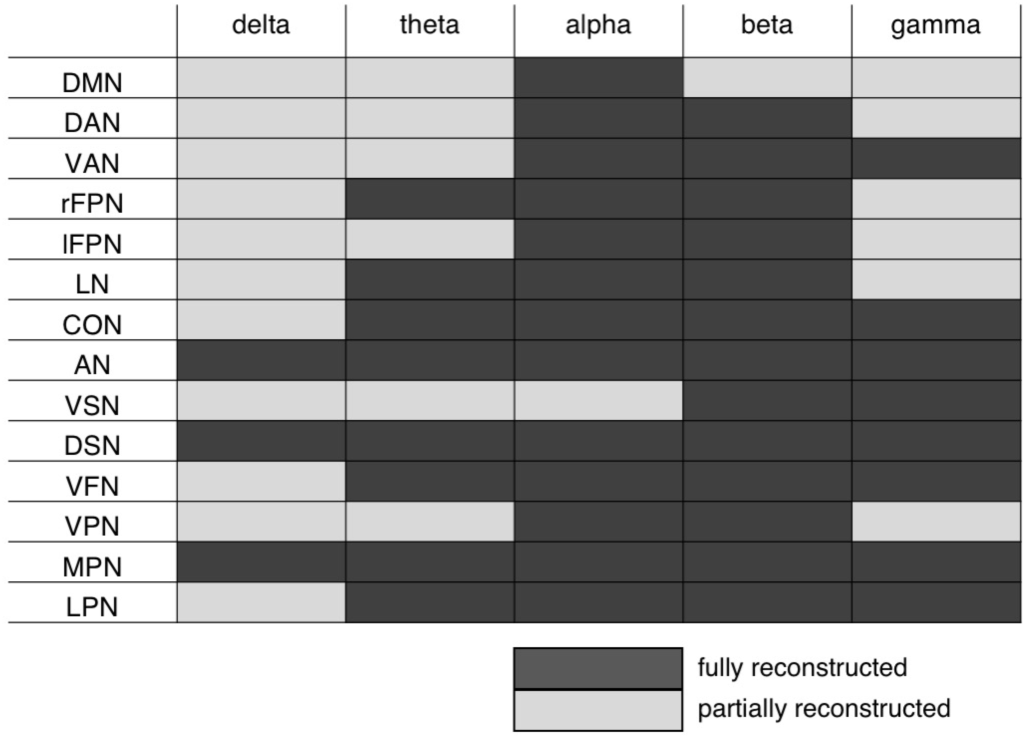
Dependence of RSN detection on the selected frequency band. We extracted band-limited power envelopes from source-space signals filtered in the following frequency bands: delta (1-4 Hz), theta (4-8 Hz), alpha (8-13 Hz), beta (13-30 Hz) and gamma (30-80 Hz). We then attempted to reconstruct 14 RSNs by tICA for each frequency band. Finally, we evaluated whether the RSN maps could be fully or only partially reconstructed.

**Supplementary Fig. 8.**
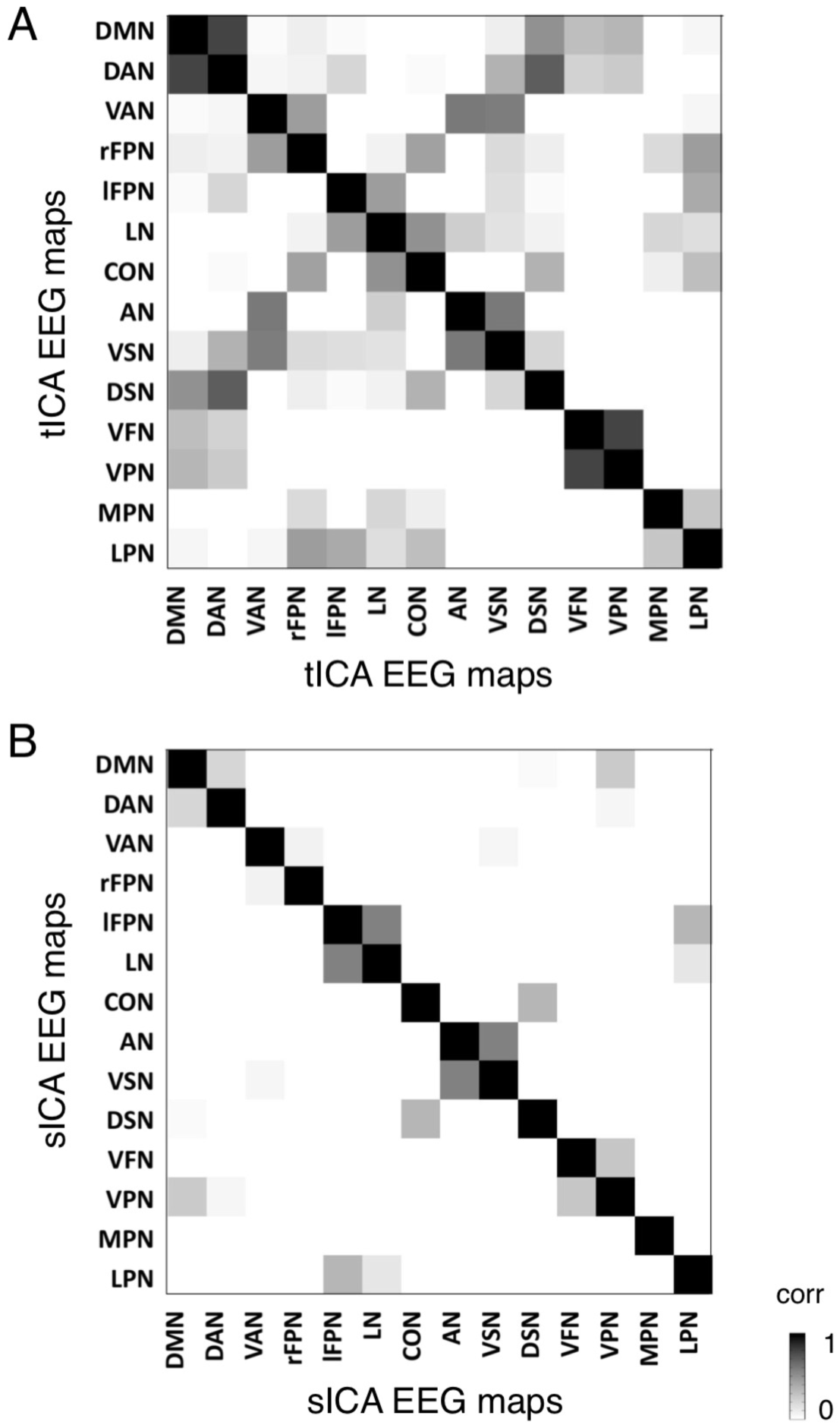
Comparison of EEG-RSN maps reconstructed using tICA and sICA. (A) Spatial correlation between EEG-RSNs detected with tICA; (B) Spatial correlation between EEG-RSNs detected with sICA. Lower non-diagonal values indicate that the spatial patterns of the different RSNs are more distinct.

**Supplementary Table 1.**
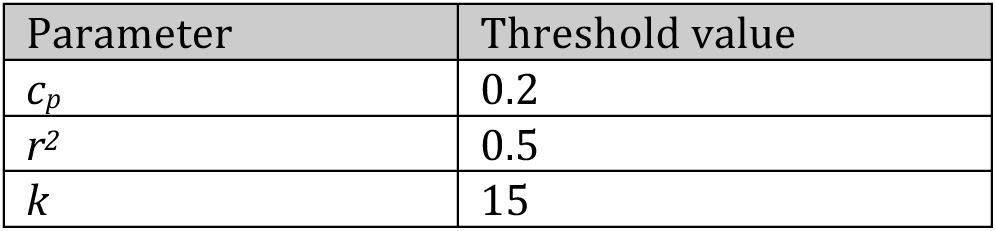
Thresholds used for the automated detection of artifactual ICs. These were set in accordance with previous EEG/MEG studies (de Pasquale et al., 2010; Mantini et al., 2009).

**Supplementary Table 2.** Conductivity values of different tissues used for the calculation of the head model. The conductivity values associated with the tissue classes were extracted from relevant literature (Haueisen et al., 1997).

| Tissue name | Conductivity (S/m) |
| --- | --- |
| Skin | 0.4348 |
| compact bone | 0.0063 |
| spongy bone | 0.0400 |
| CSF | 1.5385 |
| cortical gray matter | 0.3333 |
| cerebellar gray matter | 0.2564 |
| cortical white matter | 0.1429 |
| cerebellar white matter | 0.1099 |
| brainstem | 0.1538 |
| eyes | 0.5000 |
| muscle | 0.1000 |
| fat | 0.0400 |

## References

Akalin Acar, Z., Acar, C.E., Makeig, S., 2016. Simultaneous head tissue conductivity and EEG source location estimation. Neuroimage 124, 168–180.

Baker, A.P., Brookes, M.J., Rezek, I.A., Smith, S.M., Behrens, T., Smith, P.J.P., Woolrich, M., 2014. Fast transient networks in spontaneous human brain activity. Elife 3.

Besl, P.J., Mckay, N.D., 1992. A Method for Registration of 3-D Shapes. Ieee Transactions on Pattern Analysis and Machine Intelligence 14, 239–256.

Brookes, M.J., Woolrich, M., Luckhoo, H., Price, D., Hale, J.R., Stephenson, M.C., Barnes, G.R., Smith, S.M., Morris, P.G., 2011. Investigating the electrophysiological basis of resting state networks using magnetoencephalography. Proc Natl Acad Sci U S A 108, 16783–16788.

Brookes, M.J., Woolrich, M.W., Barnes, G.R., 2012. Measuring functional connectivity in MEG: a multivariate approach insensitive to linear source leakage. Neuroimage 63, 910–920.

Calhoun, V.D., Adali, T., Pearlson, G.D., Pekar, J.J., 2001. Spatial and temporal independent component analysis of functional MRI data containing a pair of task-related waveforms. Human Brain Mapping 13, 43–53.

Calhoun, V.D., Liu, J., Adali, T., 2009. A review of group ICA for fMRI data and ICA for joint inference of imaging, genetic, and ERP data. Neuroimage 45, S163–172.

Cho, J.H., Vorwerk, J., Wolters, C.H., Knosche, T.R., 2015. Influence of the head model on EEG and MEG source connectivity analyses. Neuroimage 110, 60–77.

Dale, A.M., Liu, A.K., Fischl, B.R., Buckner, R.L., Belliveau, J.W., Lewine, J.D., Halgren, E., 2000. Dynamic statistical parametric mapping: Combining fMRI and MEG for high-resolution imaging of cortical activity. Neuron 26, 55–67.

de Pasquale, F., Della Penna, S., Snyder, A.Z., Lewis, C., Mantini, D., Marzetti, L., Belardinelli, P., Ciancetta, L., Pizzella, V., Romani, G.L., Corbetta, M., 2010. Temporal dynamics of spontaneous MEG activity in brain networks. Proc Natl Acad Sci U S A 107, 6040–6045.

de Pasquale, F., Della Penna, S., Snyder, A.Z., Marzetti, L., Pizzella, V., Romani, G.L., Corbetta, M., 2012. A cortical core for dynamic integration of functional networks in the resting human brain. Neuron 74, 753–764.

Deco, G., Jirsa, V.K., McIntosh, A.R., 2011. Emerging concepts for the dynamical organization of resting-state activity in the brain. Nature Reviews Neuroscience 12, 43–56.

Fiederer, L.D., Vorwerk, J., Lucka, F., Dannhauer, M., Yang, S., Dumpelmann, M., Schulze-Bonhage, A., Aertsen, A., Speck, O., Wolters, C.H., Ball, T., 2015. The role of blood vessels in high-resolution volume conductor head modeling of EEG. Neuroimage 128, 193–208.

Fox, M.D., Raichle, M.E., 2007. Spontaneous fluctuations in brain activity observed with functional magnetic resonance imaging. Nat.Rev.Neurosci. 8, 700–711.

Friston, K., 2002. Beyond phrenology: what can neuroimaging tell us about distributed circuitry? Annu Rev Neurosci 25, 221–250.

Friston, K.J., 2011. Functional and effective connectivity: a review. Brain Connect. 1, 13–36.

Fuchs, M., Kastner, J., Wagner, M., Hawes, S., Ebersole, J.S., 2002. A standardized boundary element method volume conductor model. Clin Neurophysiol 113, 702–712.

Ganzetti, M., Mantini, D., 2013. Functional connectivity and oscillatory neuronal activity in the resting human brain. Neuroscience 240, 297–309.

Gillebert, C.R., Mantini, D., 2013. Functional connectivity in the normal and injured brain. Neuroscientist. 19, 509–522.

Haueisen, J., Ramon, C., Eiselt, M., Brauer, H., Nowak, H., 1997. Influence of tissue resistivities on neuromagnetic fields and electric potentials studied with a finite element model of the head. IEEE Trans Biomed Eng 44, 727–735.

Hillebrand, A., Barnes, G.R., Bosboom, J.L., Berendse, H.W., Stam, C.J., 2012. Frequency-dependent functional connectivity within resting-state networks: an atlas-based MEG beamformer solution. Neuroimage 59, 3909–3921.

Himberg, J., Hyvarinen, A., 2003. ICASSO: Software for investigating the reliability of ICA estimates by clustering and visualization. 2003 Ieee Xiii Workshop on Neural Networks for Signal Processing - Nnsp’03, 259–268.

Hipp, J.F., Engel, A.K., Siegel, M., 2011. Oscillatory synchronization in large-scale cortical networks predicts perception. Neuron 69, 387–396.

Hipp, J.F., Hawellek, D.J., Corbetta, M., Siegel, M., Engel, A.K., 2012. Large-scale cortical correlation structure of spontaneous oscillatory activity. Nature Neuroscience 15, 884–890.

Holdefer, R.N., Sadleir, R., Russell, M.J., 2006. Predicted current densities in the brain during transcranial electrical stimulation. Clin Neurophysiol 117, 1388–1397.

Iacono, M.I., Neufeld, E., Akinnagbe, E., Bower, K., Wolf, J., Vogiatzis Oikonomidis, I., Sharma, D., Lloyd, B., Wilm, B.J., Wyss, M., Pruessmann, K.P., Jakab, A., Makris, N., Cohen, E.D., Kuster, N., Kainz, W., Angelone, L.M., 2015. MIDA: A Multimodal Imaging-Based Detailed Anatomical Model of the Human Head and Neck. PLoS One 10, e0124126.

Johnston, M.V., 2004. Clinical disorders of brain plasticity. Brain Dev 26, 73–80.

Lew, S., Wolters, C.H., Anwander, A., Makeig, S., MacLeod, R.S., 2009. Improved EEG Source Analysis Using Low-Resolution Conductivity Estimation in a Four-Compartment Finite Element Head Model. Human Brain Mapping 30, 2862–2878.

Li, Y.O., Adali, T., Calhoun, V.D., 2007. Estimating the number of independent components for functional magnetic resonance imaging data. Human Brain Mapping 28, 1251–1266.

Lin, F.H., Witzel, T., Ahlfors, S.P., Stufflebeam, S.M., Belliveau, J.W., Hamalainen, M.S., 2006. Assessing and improving the spatial accuracy in MEG source localization by depth-weighted minimum-norm estimates. Neuroimage 31, 160–171.

Liu, Q., Balsters, J.H., Baechinger, M., van der Groen, O., Wenderoth, N., Mantini, D., 2015. Estimating a neutral reference for electroencephalographic recordings: the importance of using a high-density montage and a realistic head model. Journal of Neural Engineering 12, 056012.

Mantini, D., Corbetta, M., Perrucci, M.G., Romani, G.L., Del Gratta, C., 2009. Large-scale brain networks account for sustained and transient activity during target detection. Neuroimage 44, 265–274.

Mantini, D., Corbetta, M., Romani, G.L., Orban, G.A., Vanduffel, W., 2013. Evolutionarily novel functional networks in the human brain? Journal of Neuroscience 33, 3259–3275.

Mantini, D., Della Penna, S., Marzetti, L., de Pasquale, F., Pizzella, V., Corbetta, M., Romani, G.L., 2011. A signal-processing pipeline for magnetoencephalography resting-state networks. Brain Connect 1, 49–59.

Mantini, D., Franciotti, R., Romani, G.L., Pizzella, V., 2008. Improving MEG source localizations: an automated method for complete artifact removal based on independent component analysis. Neuroimage 40, 160–173.

Mantini, D., Perrucci, M.G., Del Gratta, C., Romani, G.L., Corbetta, M., 2007. Electrophysiological signatures of resting state networks in the human brain. Proc Natl Acad Sci U S A 104, 13170–13175.

Marino, M.; Liu, Q.; Brem, S.; Wenderoth, N.; Mantini, D., 2016. Automated detection and labeling of high-density EEG electrodes from structural MR images. Journal of Neural Engineering 13.

Marzetti, L., Della Penna, S., Snyder, A.Z., Pizzella, V., Nolte, G., de Pasquale, F., Romani, G.L., Corbetta, M., 2013. Frequency specific interactions of MEG resting state activity within and across brain networks as revealed by the multivariate interaction measure. Neuroimage 79, 172–183.

McKeown, M.J., Makeig, S., Brown, G.G., Jung, T.P., Kindermann, S.S., Bell, A.J., Sejnowski, T.J., 1998. Analysis of fMRI data by blind separation into independent spatial components. Human Brain Mapping 6, 160–188.

Michel, C.M., Murray, M.M., Lantz, G., Gonzalez, S., Spinelli, L., Grave de Peralta, R., 2004. EEG source imaging. Clin Neurophysiol 115, 2195–2222.

Palva, J.M., Palva, S., 2012. Infra-slow fluctuations in electrophysiological recordings, blood-oxygenation-level-dependent signals, and psychophysical time series. Neuroimage 62, 2201–2211.

Pascual-Marqui, R.D., Lehmann, D., Koukkou, M., Kochi, K., Anderer, P., Saletu, B., Tanaka, H., Hirata, K., John, E.R., Prichep, L., Biscay-Lirio, R., Kinoshita, T., 2011. Assessing interactions in the brain with exact low-resolution electromagnetic tomography. Philos Trans A Math Phys Eng Sci 369, 3768–3784.

Pfurtscheller, G., Lopes da Silva, F.H., 1999. Event-related EEG/MEG synchronization and desynchronization: basic principles. Clin.Neurophysiol. 110, 1842–1857.

Ramon, C., Schimpf, P.H., Haueisen, J., 2006. Influence of head models on EEG simulations and inverse source localizations. Biomed Eng Online 5, 10.

Russell, G.S., Jeffrey Eriksen, K., Poolman, P., Luu, P., Tucker, D.M., 2005. Geodesic photogrammetry for localizing sensor positions in dense-array EEG. Clin Neurophysiol 116, 1130–1140.

Schaul, N., 1998. The fundamental neural mechanisms of electroencephalography. Electroencephalogr Clin Neurophysiol 106, 101–107.

Slutzky, M.W., Jordan, L.R., Krieg, T., Chen, M., Mogul, D.J., Miller, L.E., 2010. Optimal spacing of surface electrode arrays for brain-machine interface applications. Journal of Neural Engineering 7, 26004.

Smit, D.J., Stam, C.J., Posthuma, D., Boomsma, D.I., de Geus, E.J., 2008. Heritability of “small-world” networks in the brain: a graph theoretical analysis of resting-state EEG functional connectivity. Human Brain Mapping 29, 1368–1378.

Smith, S.M., Miller, K.L., Moeller, S., Xu, J., Auerbach, E.J., Woolrich, M.W., Beckmann, C.F., Jenkinson, M., Andersson, J., Glasser, M.F., Van Essen, D.C., Feinberg, D.A., Yacoub, E.S., Ugurbil, K., 2012. Temporally-independent functional modes of spontaneous brain activity. Proc Natl Acad Sci U S A 109, 3131–3136.

Smith, S.M., Nichols, T.E., 2009. Threshold-free cluster enhancement: addressing problems of smoothing, threshold dependence and localisation in cluster inference. Neuroimage 44, 83–98.

Song, J., Davey, C., Poulsen, C., Luu, P., Turovets, S., Anderson, E., Li, K., Tucker, D., 2015. EEG source localization: Sensor density and head surface coverage. J Neurosci Methods 256, 9–21.

Van de Ville, D., Britz, J., Michel, C.M., 2010. EEG microstate sequences in healthy humans at rest reveal scale-free dynamics. Proc Natl Acad Sci U S A 107, 18179–18184.

Van Hoey, G., Vanrumste, B., D’Have, M., Van de Walle, R., Lemahieu, I., Boon, P., 2000. Influence of measurement noise and electrode mislocalisation on EEG dipole-source localisation. Med Biol Eng Comput 38, 287–296.

Van Uitert, R., Weinstein, D., Johnson, C., 2003. Volume currents in forward and inverse magnetoencephalographic simulations using realistic head models. Ann Biomed Eng 31, 21–31.

Varela, F., Lachaux, J.P., Rodriguez, E., Martinerie, J., 2001. The brainweb: phase synchronization and large-scale integration. Nat.Rev.Neurosci. 2, 229–239.

Wagner, S., Rampersad, S.M., Aydin, U., Vorwerk, J., Oostendorp, T.F., Neuling, T., Herrmann, C.S., Stegeman, D.F., Wolters, C.H., 2014. Investigation of tDCS volume conduction effects in a highly realistic head model. Journal of Neural Engineering 11, 016002.

Wang, Y.H., Gotman, J., 2001. The influence of electrode location errors on EEG dipole source localization with a realistic head model. Clinical Neurophysiology 112, 1777–1780.

Wolters, C.H., Anwander, A., Tricoche, X., Weinstein, D., Koch, M.A., MacLeod, R.S., 2006. Influence of tissue conductivity anisotropy on EEG/MEG field and return current computation in a realistic head model: a simulation and visualization study using high-resolution finite element modeling. Neuroimage 30, 813–826.

Wolters, C.H., Grasedyck, L., Hackbusch, W., 2004. Efficient computation of lead field bases and influence matrix for the FEM-based EEG and MEG inverse problem. Inverse Problems 20, 1099–1116.

Yuan, H., Ding, L., Zhu, M., Zotev, V., Phillips, R., Bodurka, J., 2016. Reconstructing Large-Scale Brain Resting-State Networks from High-Resolution EEG: Spatial and Temporal Comparisons with fMRI. Brain Connect 6, 122–135.

